# Computational fluid dynamics provide evidence for a compensatory suction feeding like effect in the predatory strike of dragonfly larvae

**DOI:** 10.1101/2021.06.15.448319

**Authors:** Alexander Koehnsen, Martin Brede, Stanislav N Gorb, Sebastian Büsse

## Abstract

Most fast-moving aquatic predators face the challenge of bow wave formation. Water in front of the predator alarms or even displaces the prey. To mitigate the formation of such a bow wave, a strategy aiming at pressure reduction via suction has evolved convergently in several animal groups: compensatory suction feeding. The aquatic larvae of dragonflies and damselflies (Insecta: Odonata) are likely to face this challenge as well. They capture prey underwater using a fast-moving raptorial appendage, the so-called prehensile labial mask. Within dragonflies (Odonata: Anisoptera) two basic shapes of the prehensile labial mask have evolved, with an either flat and slender or concave distal segment. While the former is a pure grasping device, the latter is also capable of scooping up smaller prey and retaining it inside the cavity by arrays of bristle-like structures. The hydrodynamics of the prehensile labial mask was previously unknown. We used computational fluid dynamic (CFD) simulations of the distal segment of the mask, to investigate for the first time how different shapes of the mask impact their function. Our results suggest that both shapes are highly streamlined and generate a low-pressure area, likely leading to an effect analogous to compensatory suction feeding. This presents a vivid example of how convergent evolution enables very different animal groups to successfully deal with the challenges of their environment. It may also be an interesting concept for technical application in small scale grasping devices; e.g. for simple sampling mechanisms in small-sized autonomous underwater vehicles (μAUVs).

## Introduction

“Everything was an artefact of its function. That was what made evolution so gorgeous.” (Corey 2014 p. 64). In this study, we explored the relationship between the form of a prey capturing device in dragonfly larvae and their hydrodynamic implications - which in the end, represents a paragon for convergent evolution.

Most aquatic predators face a challenge when it comes to prey capturing: fast underwater motion creates a bow wave in front of the predators head, displacing or alarming prey before capturing (Lauder and Prendergast 1992; Alfaro 2002; Holzman and Wainwright 2009).

The hydrodynamics of other underwater feeding and/or prey-capturing mechanisms have been studied primarily in vertebrates (Wainwright et al. 2015): for example snapping turtles (Lauder and Prendergast 1992), amphibious snakes (Alfaro, 2002) and especially fish (e.g. Edmonds et al. 2001; Yaniv et al. 2014; Van Wassenbergh 2015). Several strategies have evolved in these groups to mitigate bow wave formation. For example, many fish species rely on suction feeding (Muller and Osse 1984; Edmonds et al. 2001; Holzman and Wainwright 2009; Yaniv et al. 2014; Van Wassenbergh 2015) and some aquatic snakes use a lateral head movement (Young 1991). However, frogs (Carroño and Nishikawa 2010), snapping turtles (Lauder and Prendergast 1992) and some snakes project their head directly towards the prey (Alfaro 2002) potentially causing bow wave formation. This problem is overcome in these groups by a mechanism called compensatory suction feeding. For example, snapping turtles quickly open their mouth while projecting their head towards a target (Lauder and Prendergast 1992; Summers et al. 1998). By flooding the oral cavity, water is drawn from the bulk which is amassing in front of the accelerating head (Lauder and Prendergast 1992; Summers et al. 1998). The mechanism thereby avoids the formation of a bow wave, allowing for successful prey capturing (Lauder and Prendergast 1992; Summers et al. 1998). As this issue gets more difficult with smaller scale and smaller prey items/particles (China and Holzman 2014), it may be a factor influencing the prey capturing in smaller invertebrates as well. In aquatic arthropods, the hydrodynamics of the predatory strike of mantis shrimps and snapping shrimps have been investigated (McHenry et al. 2016; Koukouvinis et al. 2017), but these studies did not focus on the interaction between predator and prey item so far.

In the larvae of damselflies and dragonflies (Insecta: Odonata), a unique prey capturing mechanism has evolved: one mouthpart, the labium, is modified into a prehensile mask (PLM) (see figure 1); capable of rapid projection for prey capturing (Corbet, 1957; Pritchard, 1965; Olesen, 1979; Büsse et al. 2021). The prey spectra between larvae with these two different ecotypes of PLM differ significantly. The concave PLM can scoop up smaller prey items, which are trapped inside the premental cavity and then lead to the mouth for feeding (Pritchard 1964, 1965). The sole function of the flat PLM is grasping relatively large prey with hook-like labial palps. (Tillyard 1917; Pritchard 1965).

**Figure 1).**
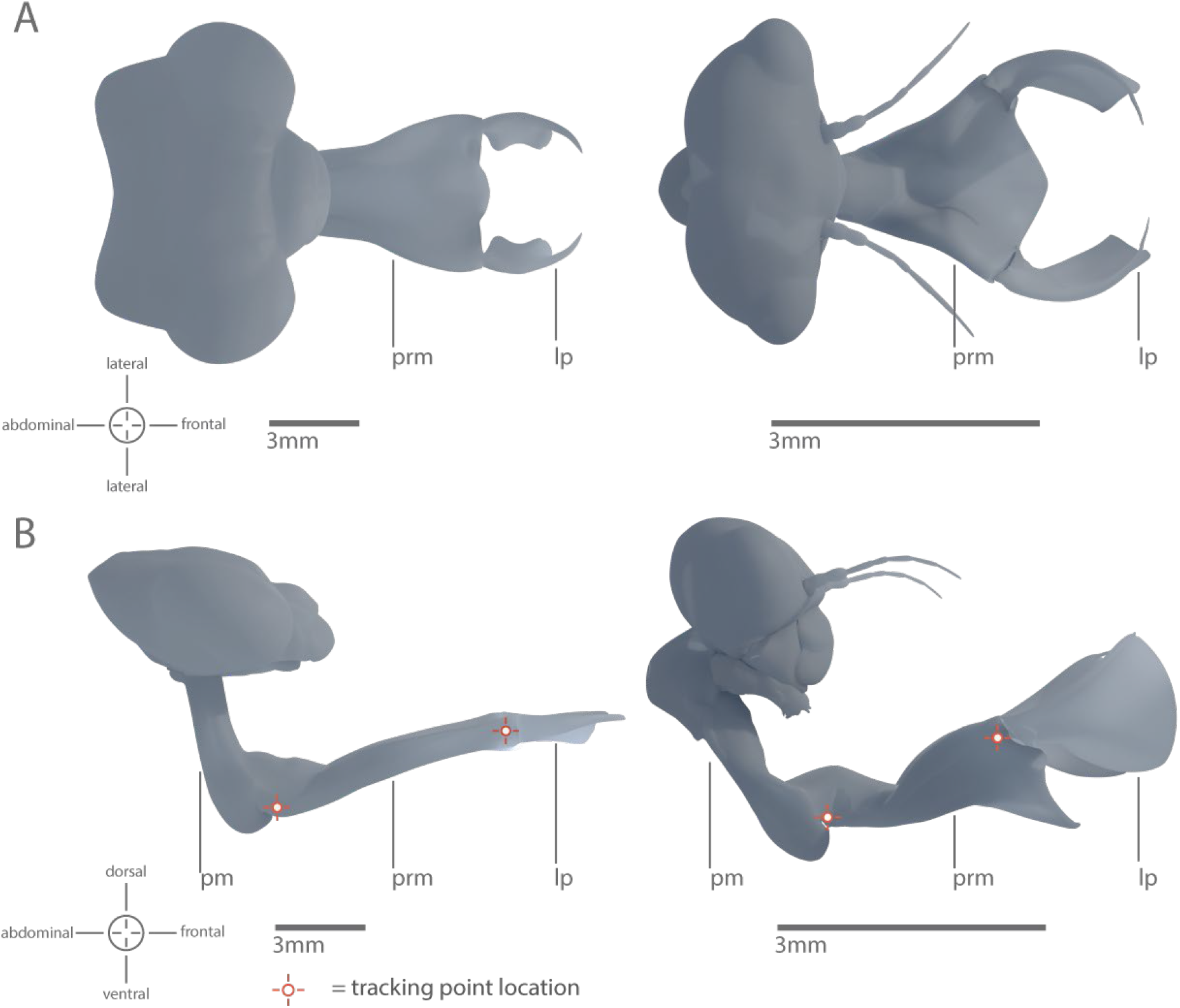
3D models of the different basic types of prehensile labial masks in dragonfly larvae (Anisoptera). Models are based on μCT scans of specimens of the aeshnid *Anax imperator* (left) which features a flat prementum for grasping large prey and the libellulid *Sympetrum* sp. (right) which features a spoon-shaped prementum also capable of scooping up smaller animals. A) dorsal view, B) lateral view; pm= postmentum, prm=prementum, lp=labial palps

The PLM in general is capable of rapid protraction (Pritchard 1965; Olesen 1979; Tanaka and Hisada 1980). Powered by a dual catapult mechanism, maximum accelerations of 20g and maximum velocities of around 1 m/s have been measured (Büsse et al. 2021). At high speeds, the influence of hydrodynamic forces on the movement becomes increasingly important, as substantial amounts of drag incur (Vogel 1997; McHenry et al. 2016;). Yet, the hydrodynamics of the PLM have never been studied previously. A few publications hypothesise about the impact of hydrodynamics on the predatory strike in their discussion (Tanaka and Hisada 1980; Quenta Herrera et al. 2018), but to the best of our knowledge, no study has put a focus on this specific topic.

In this study, we aimed to assess the two described premental shapes from a hydrodynamic point of view. Using high-speed videography, we assessed the kinematics of the prementum in detail and estimated the angle of attack during the strike for species of both ecotypes. This data was combined with 3D models obtained from micro-computed tomography (μCT) and used to generate computational fluid dynamics (CFD) simulations to resolve flow characteristics. With this approach, we intend to shed light on the question of how the two differently shaped prementa differ in their flow patterns, and how these differences might impact the respective feeding strategies. Additionally, we investigated whether the PLM features a mechanism to mitigate bow wave formation in front of the prey capturing device (Lauder and Prendergast, 1992; Alfaro 2002). This aims to further understanding how simple, small-scale underwater prey capturing/grasping devices function in general which in turn contributes to answering questions about the evolution of two different shapes within Anisoptera. Additionally, the results of this study might be interesting for potential technical applications (e.g. sample collection) by small-sized autonomous underwater vehicles (μAUVs) (Shahinpoor 1992; Kim et al. 2005; Shi et al. 2010).

## Material & Methods

### Specimens

Larvae of *Anax imperator* Leach, 1815 (Anisoptera: Aeshnidae) were collected in aquaculture ponds in Oeversee, Schleswig-Holstein, Germany. Specimens of *Sympetrum* sp. (Anisoptera: Libellulidae) were collected in a pond near Knoop, Schleswig-Holstein, Germany. All specimens were collected with permission of the *‘Landesamt für Landwirtschaft, Umwelt und ländliche Räume Schleswig-Holstein‘*. Specimens were kept in individual aquariums at 8°C to prevent premature emergence and fed with chironomid larvae (Diptera: Chironomidae). Before the experiments, the specimens were acclimatised to room temperature. Head and body length were measured as a scaling reference.

### High-speed Videography and Motion Tracking

7 predatory strikes of 5 specimens of *Anax imperator* (Odonata: Anisoptera) and 6 strikes of 5 specimens of *Sympetrum* sp. (Odonata: Anisoptera) were recorded in a small aquarium (210 × 100 × 105 mm) using chironomid larvae as prey items. Recordings were made with a Photron Fastcam SA1.1 (model 675K-M1; Photron, Pfullingen, Germany) equipped with a 105 mm/1:2.8 macro lens (Sigma, Tokyo, Japan) mounted on a Manfrotto-055 tripod with Manfrotto-410 geared head (Manfrotto, Spa, Italy). Illumination was provided using two Dedocol COOLT3 halogen spotlights (Dedotec, Berikon, Switzerland). The footage was captured with a resolution of 1024 × 1024 pixels at 5400 frames per second and saved as a 16-Bit TIFF image stack. For further analysis, the prementum was tracked using Adobe After Effects CS6 (Adobe Systems Software, San José, CA, USA) according to Koehnsen et al. (2020). One Tracker (Tracker A) was placed at the prementum-postmentum joint and a second tracker (Tracker B) was placed at the joint between prementum and labial palps (highlighted in figure 1 and supplementary figure 1).

To validate the simulations, we additionally recorded two specimens of each species (two strikes per specimen) with the previous setup. To visualize flow patterns, neutrally buoyant *Spirulina* algae were added to the water and a light source was placed behind the aquarium. Adjusting the focal plane to the centre line of the larva and using a shallow focal plane (fstop = 4.0), allowed only particles flowing around the approximate centre line of the PLM to be in focus.

### Data Analysis and Statistics

The tangential velocity was determined directly from the location data of tracker A, which is located at the centre of rotation of the prementum. To assess the rotational velocity, a vector between the two tracking points was established (AB). These points are also highlighted in figure 1 and supplementary material 1. The angle of this vector at the timepoint *t* was compared to the orientation of the vector at the beginning of the motion sequence (*t_0_*). In both cases, data were fitted using a polynomial regression with 11 degrees of freedom. Velocity was determined from the first derivative. The tangential velocity was calculated from the rotational velocity with the length of the prementum (without labial palps) as the radius *r*.

To calculate the angle of attack (*α*), we compared the orientation of the prementum (which is given by the vector AB) to a second vector indicating the direction of movement. This vector is formed by point B (the base of the prementum) and the location of point B five frames earlier, indicating the direction of motion. α is the angle between these two vectors. The location of all points used is also shown in supplementary figure S1. To account for the temporal shift between *α* and the directly measured curve, *α* has been calculated as a moving average: once with the aforementioned reference point, five frames earlier and once with a reference point five frames later. For each timepoint, we used the mean of the two values.

We extracted data from the curve at ten standardised timepoints for CFD simulations. Initially, the point of maximum total velocity (translational + tangential velocity) and the start and endpoint of motion were determined, dividing the strike into an acceleration and deceleration phase. Six equally spaced points (including starting point and point of maximum velocity) were used for the acceleration phase, and four equally spaced points (including the endpoint) were used for the deceleration phase.

All statistical analyses were performed in R Studio (v1.2.5001, the R Foundation for Statistical Computing, Vienna, Austria). For the comparison of mean velocity and *α*, a Wilcoxon Rank Sum Test was used, as the sample size was not equal. If not stated otherwise, all given values are median ± interquartile range (IQR).

The Reynolds number of the prementum was calculated for both species according to the following formula:

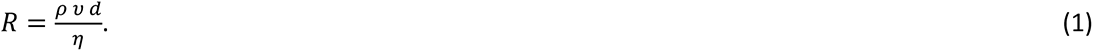

with the maximum width of the prementum (without palps) as characteristic length (*d*). The velocity (*υ*) was determined from tracking data using the total velocity at the tip. Dynamic viscosity (*η*) and density (*ρ*) were taken from literature data of freshwater at 25°C (Rumble et al. 2017) see supplementary table S2).

The drag coefficient *C_dw_* and lift coefficient *C_lw_* was calculated for both species according to the following formulas for drag coefficient and lift coefficient, respectively:

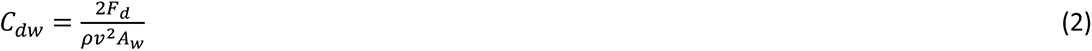

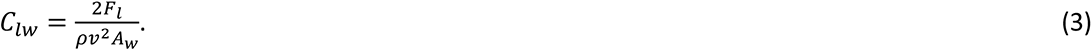

As *α* (and thereby the frontal area) of the prementum differs over the duration of the predatory strike, we used the total wetted area *A_w_* as a reference area for the calculation of C_dw_ */C_lw_*. Measurement of the reference area was based on the 3D models used for CFD simulation (supplementary figure S4). The wetted area *A_w_* was used instead of the more common projected frontal area *A_w_*, as the angle of attack, and thereby the projected frontal area would vary between simulations, complicating the interpretation of the results. *F_d_* represents the drag force obtained from the CFD simulations, while the input value for velocity (*υ*) was taken from tracking data at maximum velocity and is identical to the data used for CFD simulation. The density (*ρ*) was taken from literature data of freshwater at 25°C (supplementary table S2).

### Micro-computed tomography (μCT)

Micro-computed tomography scans were used to obtain 3D models for CFD analysis. Therefore specimens (n=1 for each species) were fixed in alcoholic Bouin solution (= Duboscq-Brasil) for tissue preservation and subsequently stored in 70% ethanol. Samples were critical-point dried before scanning (Leica EM CPD300 automated critical point dryer, Leica, Wetzlar, Germany) and scanned with a SkyScan 1172 (Bruker micro-CT, Kontich, Belgium) with high-resolution settings (40 kV, 250 μA and 0.25° rotation steps, performing a 360° scan). Segmentation and processing of the μCT-data were carried out with Amira 6.0.1 (Thermo Fischer Scientific, Waltham, USA). 3D datasets were retopologised, reoriented and rescaled using Blender (v2.8 Blender Foundation, Amsterdam, Netherlands, www.blender.org). The prementum was scaled based on the average size of all specimens (of the respective species) that were used for the high-speed video experiments.

### Scanning Electron Microscopy (SEM)

Two specimens (n=2) of *Sympetrum* sp. were prepared with extended PLM using an expendable custom-built brace (supplementary figure S5). One specimen with opened labial palps, one with closed ones. The brace was designed using blender (v2.8, Blender Foundation, Amsterdam, Netherlands, www.blender.org) and printed on a Prusa i3 Mk3 FDM 3D printer (Prusa Research s.r.o., Prague, Czech Republic) using polylactic acid filament (Prusament, Prusa Research s.r.o., Prague, Czech Republic) due to its stability in ethanol. The samples were critical-point dried (Leica EM CPD300, Leica, Wetzlar, Germany) and coated with a 10nm gold-palladium layer to (Leica Bal-TEC SCD500, Leica, Wetzlar, Germany). We mounted the samples using a rotatable sample holder (Pohl 2010) and examined them using a Hitachi TM3000 (Hitachi Ltd., Tokyo, Japan) scanning electron microscope with an acceleration voltage of 15kV. Obtained images were stitched into a single image using Affinity Photo (Serif Europe Ltd. West Brigford, UK, https://affinity.serif.com).

### Computational fluid dynamics

#### Meshing

Meshes were created using ICEM (2019 R2, Ansys Inc., Canonsburg Pa, USA). Models of the prementum of *Anax* and *Sympetrum* were imported as ‘.stl files’. We oriented the model according to the specified *α* and created a domain around the model with five times the model size as head and ten times the model size as trail region. Three times the model size was used as the clearance between model and lateral as well as top and bottom domain walls (see below for wall boundary conditions).

We created unstructured tetrahedral meshes with ^~^4.6 million elements and 5 prism layers around the model surface (maximum *y+* was below 0.69 in all simulations for *Sympetrum* and below 2 in all simulations for *Anax*). To increase mesh resolution around the model a density field was used. Doubling the element count further changed the simulation results by <3%, therefore mesh resolution was considered sufficient for our purposes (supplementary table S3). An exemplary mesh for both species is shown in supplementary figure S4.

#### Boundary conditions and solution

For simulation setup and solving, we used CFX Pre (2019 R2, Ansys Inc., Canonsburg Pa, USA) with fluid domain properties set to the values of water at 25°C. As boundary conditions, we set up the lateral as well as top and bottom walls to be free slip walls. The rear wall was defined as an opening with no ambient pressure difference. The front wall was defined as an inlet, with normal speed settings based on either the tracking data in case of the first simulated experiment or to 0.4m/s in the case of the second experiment. The model of the prementum was set up as a smooth no-slip wall. We approximated turbulence using the κε turbulence model with ANSYS default low turbulence settings. Simulations were solved using the RANS Solver Ansys CFX (2019 R2, Ansys Inc., Canonsburg PA, USA). We used RMS (root-mean-square) error as convergence criterium with a remaining error of 1×10^−4^. Increasing the convergence criteria by one order of magnitude changed the simulation result by less than 5%.

#### Simulation setup

We performed three virtual experiments using CFD Simulations: 1) We sampled 10 timepoints from each recorded strike and used the estimated *α* and flow velocity as simulation input. We used this time series of stationary simulations as a quasi-dynamic approximation to the unsteady motion of the PLM. We also neglected the rotational motion, but instead used the sum of rotational velocity (tangential velocity at the tip of the prementum) and translational velocity, as we were most interested in the area around the premental tip, where contact with the prey occurs. 2) To assess the impact of *α* on the observed flow patterns, we performed 10 stationary simulations of each prementum at a flow velocity of 0.4m/s (which is around the maximum observed flow speed), and varied *α* in ten steps between 0° and 70°. 3) We performed a scaling experiment at an *α* of 30° and 0.4 m/s flow velocity. Here we scaled down the prementum of *Anax* to the size of the prementum of *Sympetrum* (and vice versa).

## Results

### Kinematics and angle of attack

We recorded high-speed video footage of the predatory strike in *Anax* (flat-shaped) and *Sympetrum* (spoon-shaped) and calculated the translational velocity (based on the motion of the base of the prementum) and the tangential velocity (based on the rotation of the tip around the base). We neither found a significant difference in maximum translational velocity between *Anax* and *Sympetrum* (Wilcox rank-sum test, p=0.35) nor maximum tangential velocity (Wilcox rank-sum test, p=0.13). We calculated the angle of attack (*α*) based on tracking data: the angle of the prementum relative to the flow direction. Figure 2 shows the *α* (a) and the total velocity (translational + tangential velocity) (b) for both species at 10 sample points. The results suggest that the median *α* is higher in *Sympetrum* at every timepoint, yet this difference is only statistically significant at three timepoints (TP_1_ p=0.038, TP_3_ p=0.003, TP_4_ p=0.003 Wilcox rank-sum test). The velocity at each timepoint varied widely between individuals (Fig 2 B) and we found no statistically significant difference in strike velocity between both species at any timepoint (0.2 > p < 0.95, Wilcox rank-sum test). The median maximum velocity is around 0.45 +- 0.32 m/s in *Anax* and 0.43 +- 0.30 m/s in *Sympetrum*. Figure 2 A+B suggests that *α* decreases with an increased velocity and increases again towards the end of the strike as the PLM slows down.

**Figure 2).**
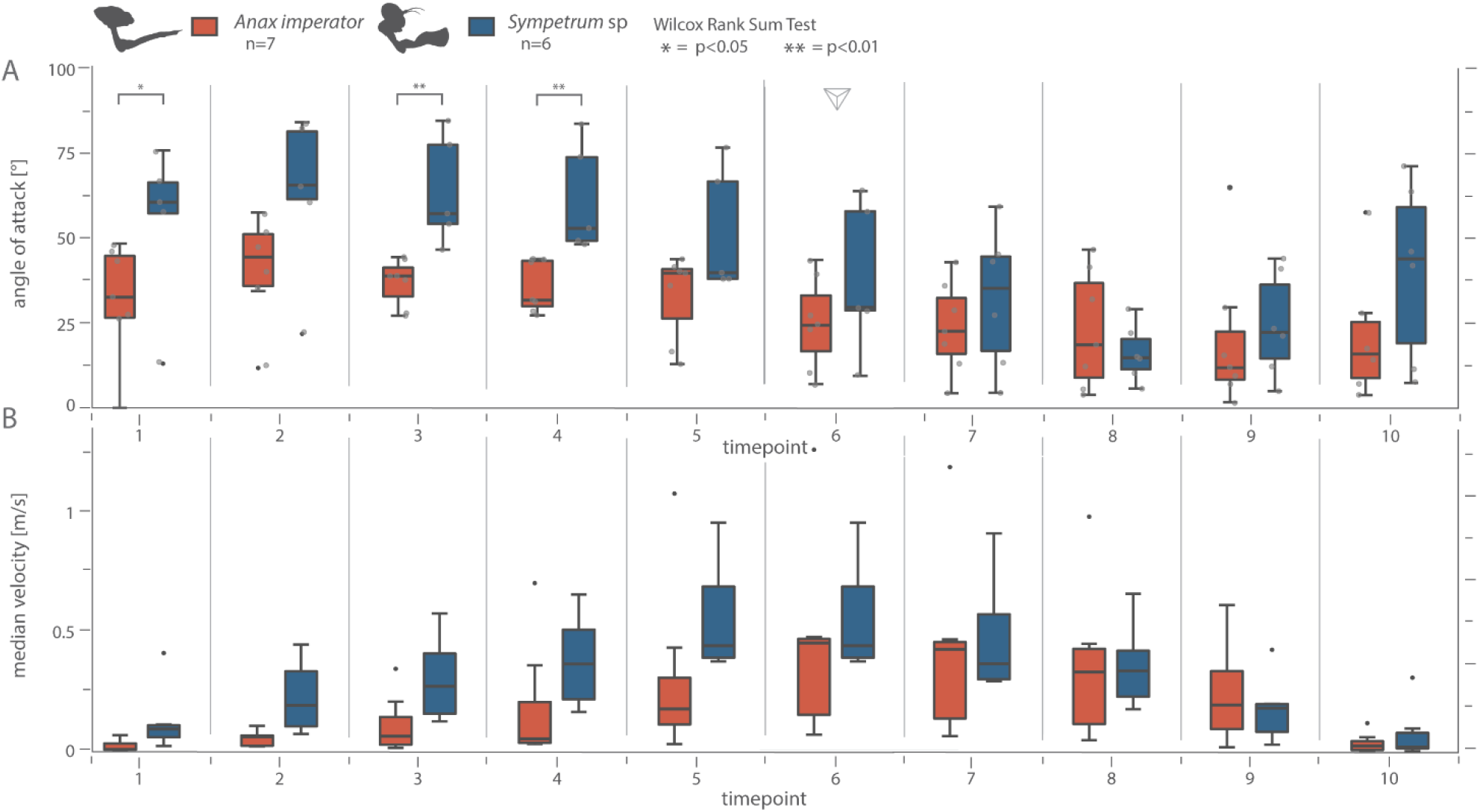
The angle of attack and total velocity of the prementum over the course of the predatory strike. A) Angle of attack (α) of the prementum of each species for 10 timepoints extracted from strike data. We see a significantly higher α at three timepoints. (Wilcox Rank Sum Test, p<0.05. *Sympetrum* n=6, *Anax* n=7). B) Median strike velocity for both species at each timepoint. Velocity represents the sum of translational and rotational velocity components at the tip of the prementum. At no point, there is a significant difference in strike velocity between the two species (Wilcox Rank Sum Test, *Sympetrum* n=6, *Anax* n=7). Data extracted from High-speed footage of the predatory strike.

### Reynolds number

We estimated the Reynolds number for the prementum of both *Anax* and *Sympetrum* models based on kinematic data with the width of the prementum (without labial palps) as reference length *r*. We calculated a Reynolds number of ^~^3700 for *Anax* and ^~^1800 for *Sympetrum* (supplementary table S2).

### Computational fluid dynamics

#### Total forces on the prementum

We performed a total of 46 computational fluid dynamics (CFD) simulations of the prementum of both species. We performed stationary simulations for each aforementioned timepoint (Fig. 2) and calculated the total drag force (parallel to the direction of flow) and the total lift force (orthogonal to the direction of flow) acting on the prementum (cf. Force diagram in figure 3). Results are shown in figure 3 (3A data on *Anax*, 3B data on *Sympetrum*). In both species, forces increase with velocity, yet the maximum velocity and maximum drag/lift force do not necessarily occur at the same timepoint. The magnitude of forces is about twice as high in *Anax* (max. F_drag_ of 1.26 mN in *Anax* and 0.56 mN in *Sympetrum*. Max. F_lift_ of 1.38 mN in *Anax* and 0.50 mN in *Sympetrum*). This also shows that the PLM generates substantial amounts of lift. Lift forces are almost as high or even exceed (in *Anax*) the amount of occurring drag at the respective timepoint.

**Figure 3).**
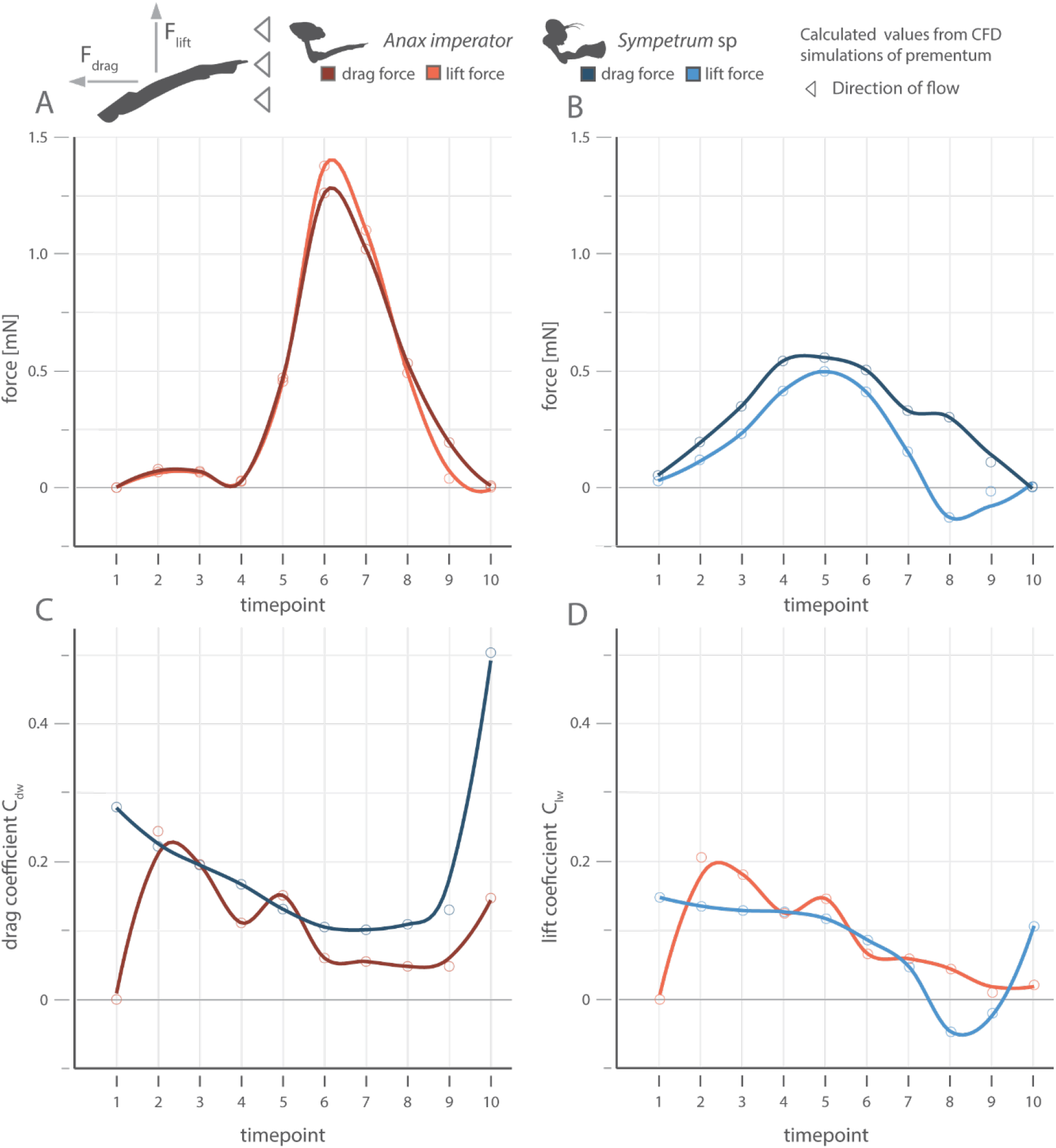
Both prehensile labial masks generate substantial amounts of lift and the drag coefficient (C_dw_) is higher in *Sympetrum*. Top left corner: Force diagram showing the direction of drag and lift force. Please note that these directions are relative to the flow directions, not the orientation of the PLM. A, B) Total drag and lift force acting on the prementum for both species at each timepoint (red *Anax*, blue *Sympetrum*, lighter colours indicate lift data). C, D) C_dw_/ C_lw_ based on the forces in A and B (same colour coding). Data points are results from CFD Simulations based on kinematic data from HS Videos (Figure 2). Lines are interpolations using a Savitzky-Golay filter (span=0.5). Please note that the drag/lift coefficient in *Anax* was zero at the first timepoint, this is an artefact due to the median velocity and forces of the prementum being too low to calculate meaningful coefficients.

#### Drag and lift coefficients

With an average prementum length of about 10 mm, the prementum of *Anax* is about three times the length of the prementum in *Sympetrum*, hence the occurring drag forces are hardly comparable. We, therefore, calculated the drag coefficient *C_dw_* and lift coefficient *C_lw_* for each timepoint (as *C_dw_* and *C_lw_* change with α). Results are shown in figures 3 C and D. The drag coefficients of both PLM start to diverge with an increased velocity (with exception of timepoint 5). During the phases with high velocity, the drag coefficient in *Anax* is about half as high as in *Sympetrum*. Drag coefficients increase again towards the end of the strike when velocity decreases and α increases. In both cases, we see a decrease in the lift coefficient (concordant with a decreasing *α*) over the course of the strike. In *Sympetrum*, we even see negative lift at timepoint 8, where *α* is lowest.

#### Pressure and flow speed profiles

We plotted contours of both the pressure (Fig. 4) and total flow speed (Fig.5) of four exemplary timepoints along the median line of the prementum. Plots for all timepoints can be found in supplementary figure S6. In both species, we see a high-pressure area below the prementum (as indicated by reddish colour) and a low-pressure region above it (as indicated by a yellowish colour). These areas are less pronounced around the prementum of *Anax* which is concurrent with its lower drag coefficient. In the steps with lower velocity (such as 3 and 9), we also see smaller pressure differences. Additional details can be inferred from the contours of the flow speed (Fig. 5). We see a higher flow speed in low-pressure areas and a decrease in high-pressure areas. In *Anax*, the flow separation is visible at higher *α*, whereas this does not occur in *Sympetrum* (see Fig. 5 step 3+5). The separation of the flow is indicated by the blue area on the dorsal side representing a slow-moving “bubble”, where the fluid has detached from the surface.

**Figure 4).**
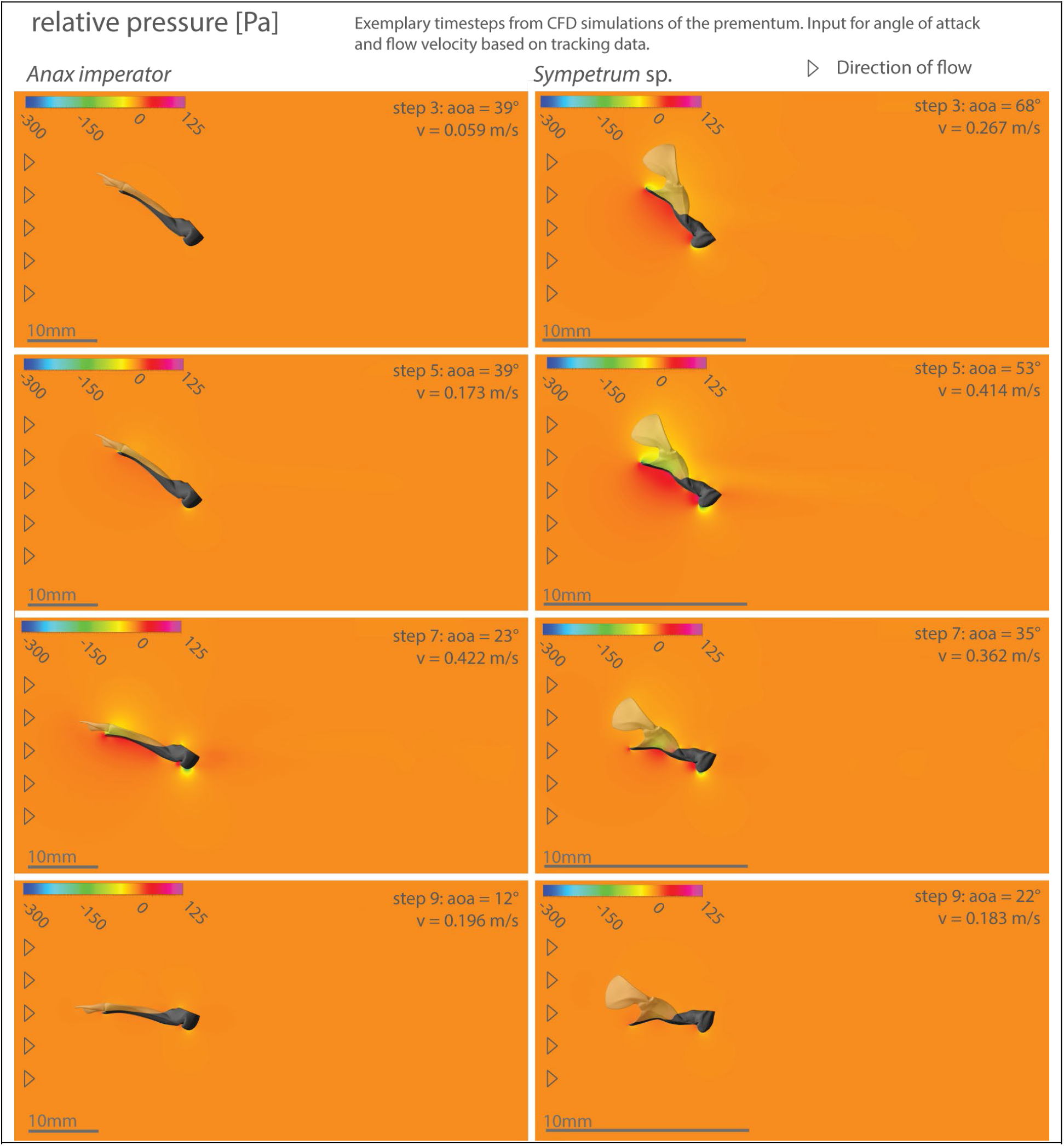
Flow characteristics change during the predatory strike I. Contours of pressure around the median line of the prementum of *Anax* l) and *Sympetrum* r). Four exemplary timepoints are shown (results from all simulations are shown in supp. figure S5). Colour indicates fluid pressure around the prementum (the colour range is identical in all images). Note the formation of low-pressure regions on the dorsal side of both prementa (yellow areas). CFD Simulations were based on kinematic data from HS Videos (Figure 2).

**Figure 5).**
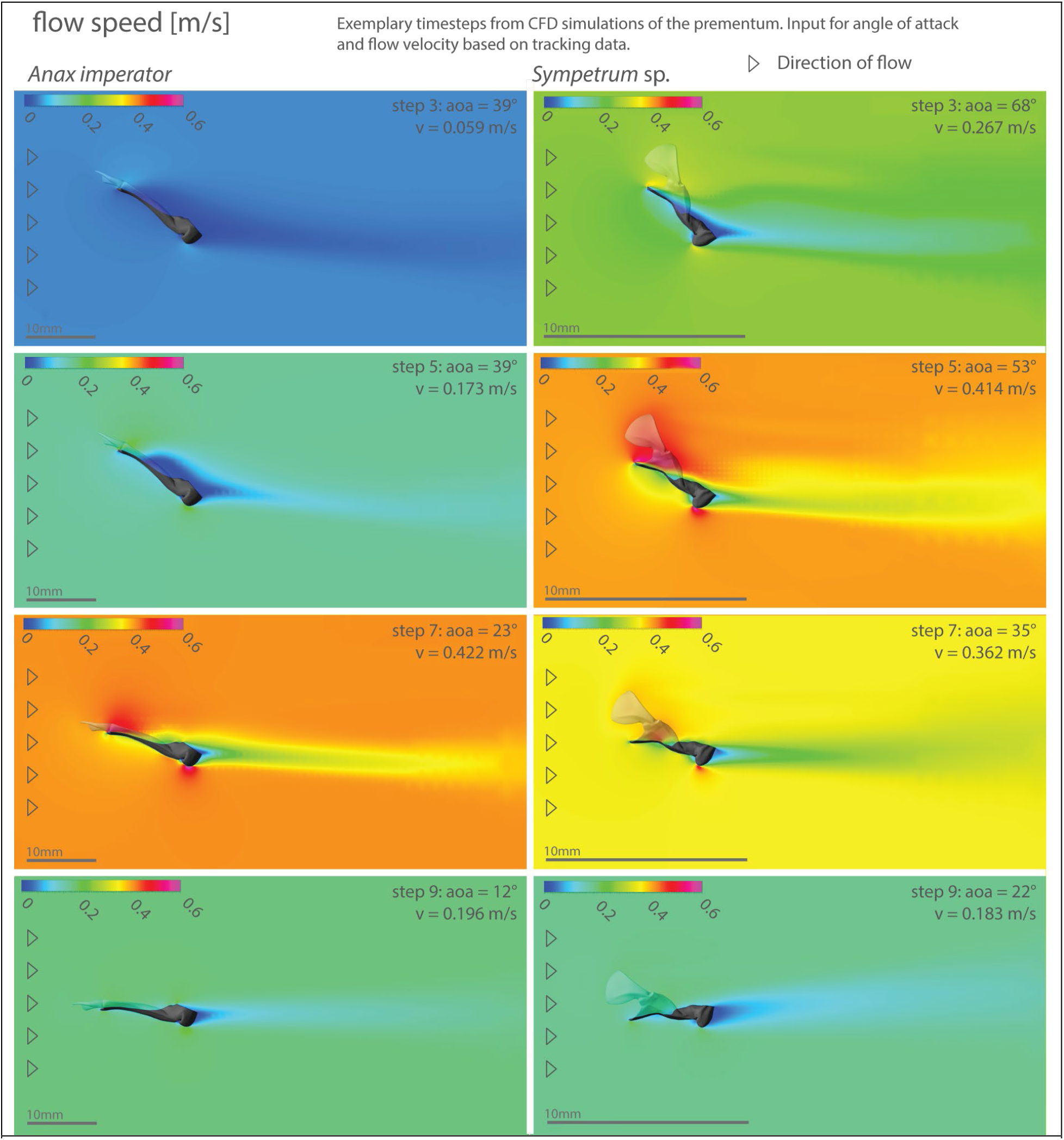
Flow characteristics change during the predatory strike II. Contours of pressure around the median line of the prementum of *Anax* l) and *Sympetrum* r). Four exemplary timepoints are shown (results from all simulations are shown in Supp. figure S6). Colour indicates fluid speed around the prementum (the colour range is identical in all images). Note the flow separation at higher α in *Anax* (Step 3,5). This does not occur in *Sympetrum*. CFD Simulations were based on kinematic data from HS Videos (Figure 2).

#### Estimating the optimal angle of attack

To assess whether the shape of the prementum of either species is optimised towards a certain angle of attack (*α*), we ran CFD simulations of the prementum at a velocity of 0.4 m/s and varied *α* within a range from 0° to 70° (Fig. 4 A). We calculated the drag (*C_dw_*) and lift (*C_lw_*) coefficients for each simulation (Fig. 4). Our results suggest that the relationship between *α* and *C_dw_/C_dw_* is different for the two PLM shapes. The lowest *C_dw_* for the prementum of *Anax* occurs at *α* of 10° (C_dw_= 0.035), while the minimum *C_dw_* for the prementum of *Sympetrum* occurs at 25° (C_dw_= 0.091). Additionally, the drag coefficient is more sensitive to changes in *α* in *Anax* than in *Sympetrum*. We found a significant increasing monotonic relationship between *α* and *C_dw_* in *Anax* (Spearman’s rank correlation, *r_s_*=0.99, p<2.2e-16), which was not present in *Sympetrum* (Spearman’s rank correlation, *r_s_* =0.51, p=0.114). While at *α* of 10°, the drag coefficient is twice as high in *Sympetrum* than in *Anax*, the drag coefficients are equal at *α*=30° and the increase continues at a steeper slope in *Anax*. The lift coefficient increases monotonically with an increasing *α* in both species (*Anax*: Spearman’s rank correlation, *r_s_*=0.75, p=0.010, *Sympetrum*: Spearman’s rank correlation, *r_s_*=0.99, p<2.2e-16). Yet while the prementum of *Anax* starts to generate positive lift (directed into the dorsal direction) between 0° and 10°, we see positive lift generation in *Sympetrum* later, between α=25° and 30°. Yet, the lift coefficient starts to decrease again in *Anax* at *α* values higher than 45°, while it continues to increase in *Sympetrum* until the second to last data point (60°).

#### Analysis of flow patterns

We visualised contours of both the pressure and total flow speed around the median line of the respective prementum. Results of four exemplary timepoints are shown in figure 7 (pressure) and figure 8 (flow speed). Results of all simulations are shown in supplementary material S7. The shape and size of low- and high-pressure regions are dependent on *α* and the species. In *Anax*, the pressure distribution is neutral at *α*=10° and we see the aforementioned pattern from *α*=20° onwards (which corresponds with a positive lift coefficient, as shown in figure 6). At high *α*, the growth curve of the lift coefficient flattens and decreases again beyond *α*=50°. At this time, the ventral side of the prementum mask is almost perpendicular to the flow direction, incurring high amounts of pressure drag. This also happens in *Sympetrum*, but at higher α (> 60°) and on a lesser scale (due to the pressure difference between the high-pressure region and the surrounding fluid). In *Sympetrum*, we see that the tip of the prementum (also known as ligula, which is tilted downwards) causes fluid to amass in the premental cavity and form a high-pressure region. This corresponds to the higher drag coefficients (Fig. 6). At *α*=30° and beyond, we see a ventral high-pressure region and a dorsal low-pressure region, the latter is formed inside the premental cavity and both contribute to lift generation. The flow speed is visualised in figure 8. We can see the flow separation on the dorsal side of the *Anax* prementum. The point of separation is already visible at *α*=30° and moves towards the tip with an increasing α. Separation of the flow is indicated by the slowly flowing region (blue). Flow separation occurs in *Sympetrum* as well, yet to a lesser degree and only at α>=70°.

**Figure 6).**
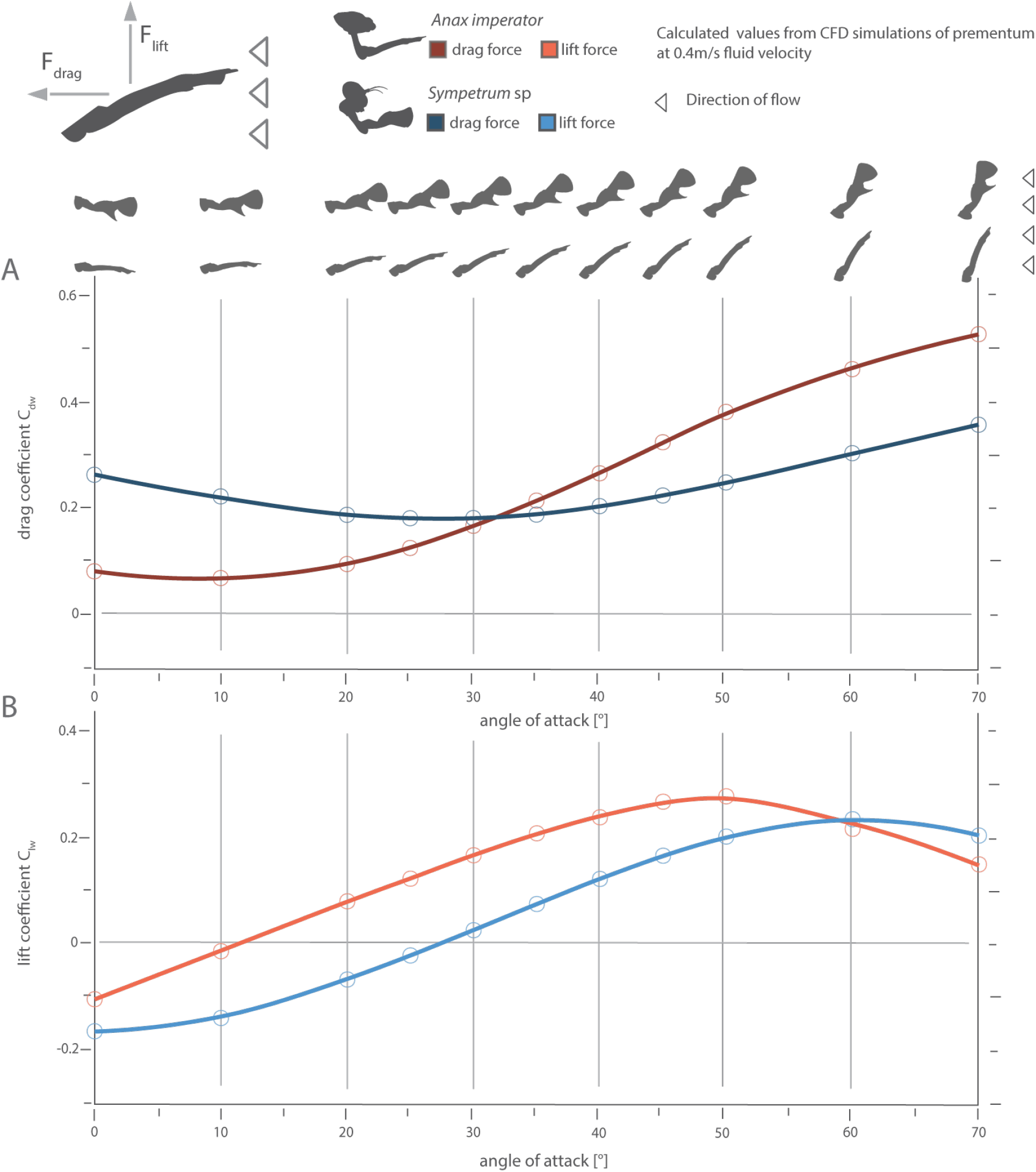
Drag and Lift Coefficient vary with the angle of attack (α). Top left corner: force diagram showing the direction of drag and lift force. Please note that these directions are relative to the flow directions. A) drag coefficient of the prementum of *Anax* (red) and *Sympetrum* (blue) vs. α. Icons at the top of the plot show orientation of the prementum to flow (flow direction right to left). B) lift coefficient of the prementum of *Anax* (red) and *Sympetrum* (blue) vs. α. In both plots: points show values from simulations, lines are interpolations using Savitzky-Golay filter. Data from CFD Simulations run at 0.4 m/s flow velocity and α was varied between 0° and 70°.

**Figure 7).**
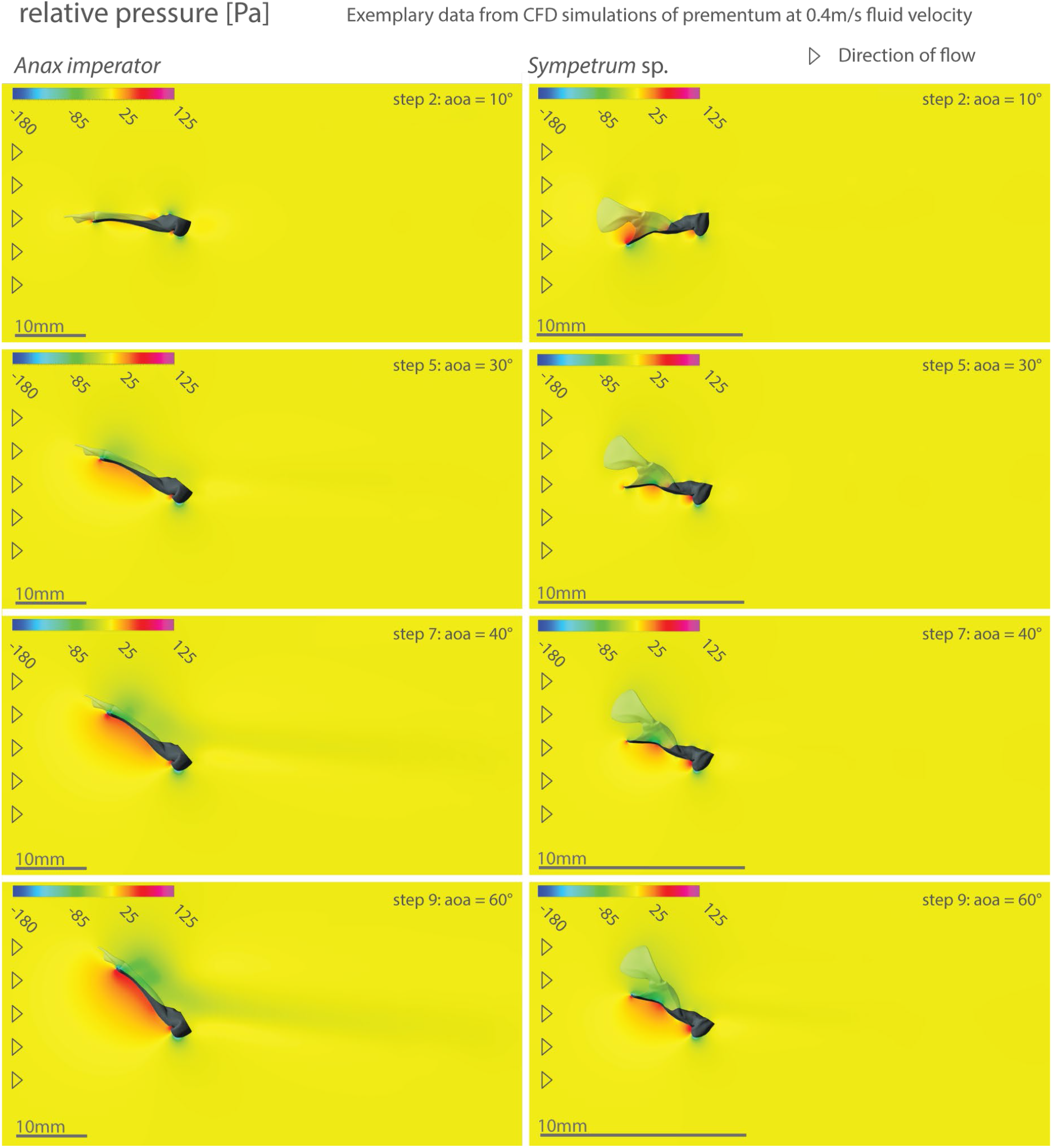
Flow characteristics change with the angle of attack (α) I. Contours of pressure around the median line of the prementum of *Anax* l) and *Sympetrum* r). Four exemplary timepoints are shown (results from all simulations are shown in Supp. figure S7). Colour indicates fluid pressure around the prementum. Note the formation of low-pressure regions on the dorsal side of both prementa at higher α. In *Sympetrum* a high-pressure area forms in the premental cavity at low α (e.g. 10°). CFD Simulations were run at 0.4m/s fluid velocity and α was varied between 0° and 70° (Figure 6).

**Figure 8.**
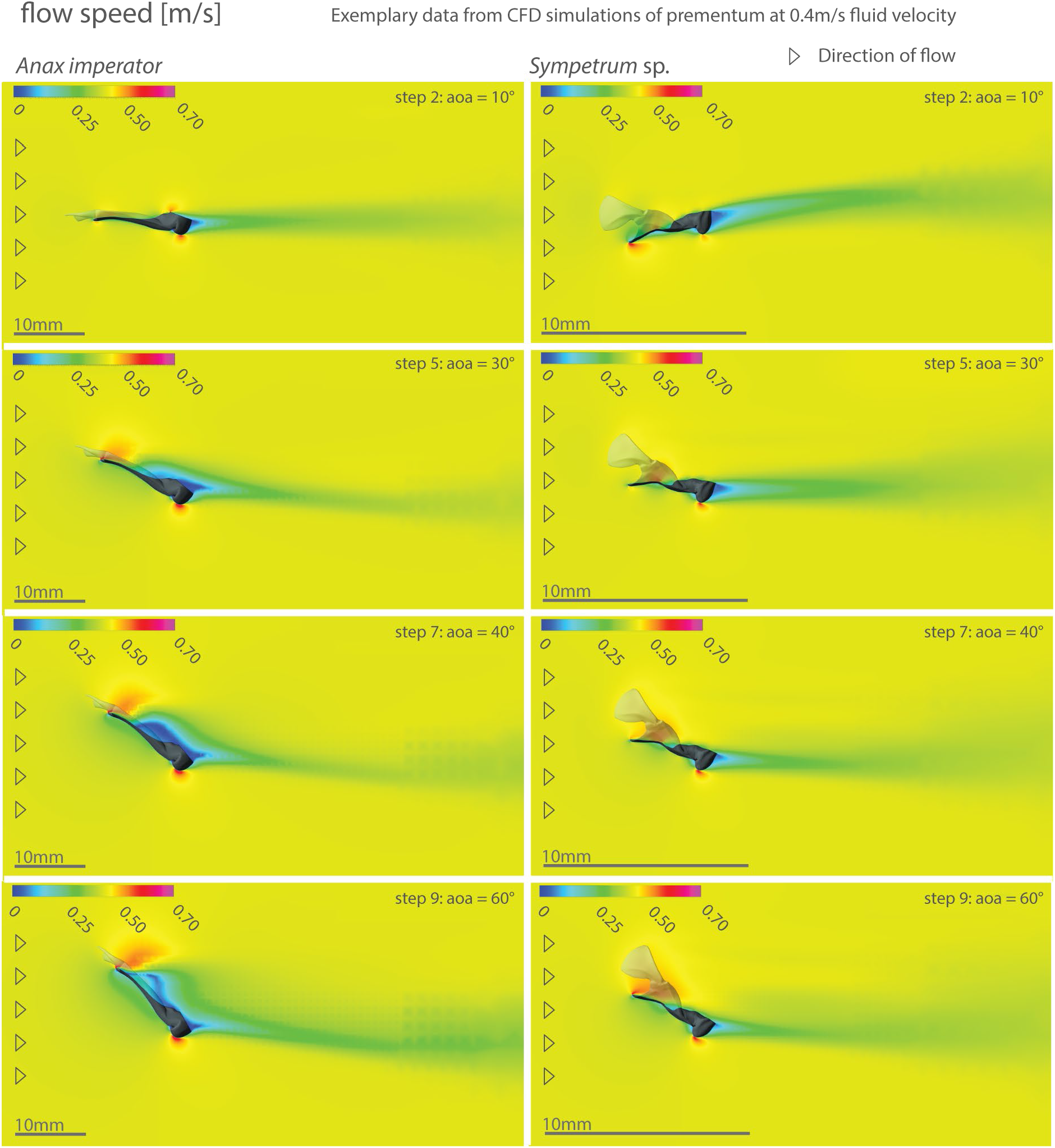
Flow characteristics change with the angle of attack (α) II. Contours of total flow speed around the median line of the prementum of *Anax* l) and *Sympetrum* r). Four exemplary timepoints are shown (Results from all simulations are shown in Supp. figure S6). Colour indicates flow speed around the prementum. Note the separation of flow occurring in the prementum of *Anax* at higher α. This does not occur in *Sympetrum*. CFD Simulations were run at 0.4m/s fluid velocity and α was varied between 0° and 70° (Figure 6).

#### Scale comparison

To assess the impact of the different Reynolds numbers on the shapes (and thereby on the comparability) of the two differently shaped PLMs, we ran an additional simulation of timepoint 5 of the previous experiment for both species. We scaled down the prementum of *Anax* to the size of the prementum of *Sympetrum* and vice versa. The results are shown in supplementary figure S8. In both species, changing the scale leads to an increase in both the drag- and the lift coefficients.

#### *Distribution of setae in* Sympetrum

Four arrays of setae cover the inside of the prementum and the labial palps (Fig. 9 A, B). Two arrays on each side of the prementum are protruded towards the centre of the premental cavity and angled forwards and downwards into the cavity. The other two arrays are located on the labial palps and protruded towards the centre of the premental cavity. When the labial palps are closed, the setae cover the premental cavity completely (Fig. 9 B), but they leave an opening during the strike when the labial palps are open (Fig. 9 A).

**Figure 9).**
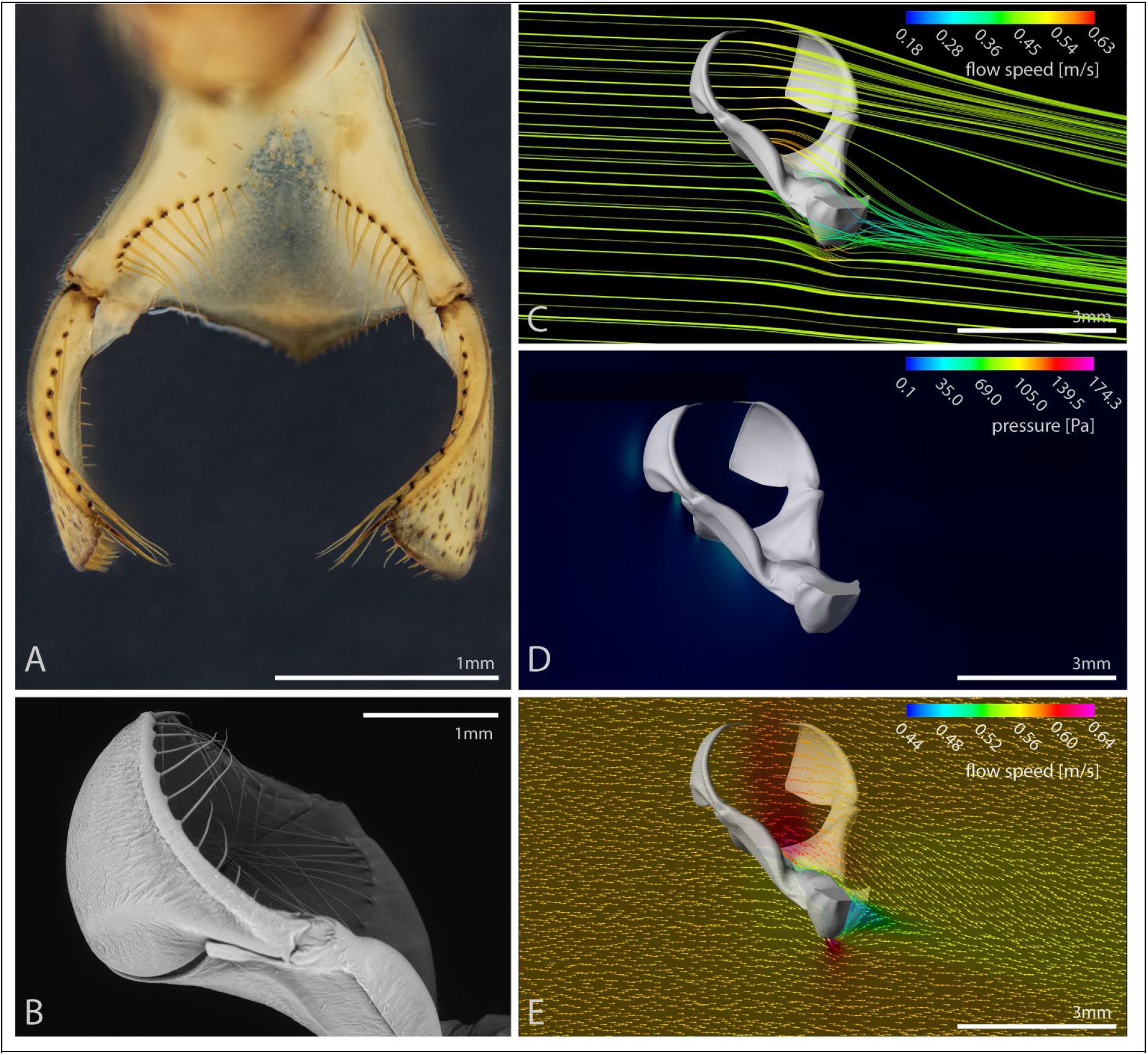
Flow is directed to a low-pressure area where setae are located. A) Photograph of the PLM of *Sympetrum*. Four arrays of setae can be identified, one on each side of the premental cavity, one on each labial palp. Note that they do not completely cover the opening, but only the premental cavity. B) SEM Micrograph of the Prementum with closed labial palps. Note that setae now cover the entire premental cavity. C) Streamlines show that flow is directed downwards above the prementum, very likely into the setae. D) 3D Heatmap showing the pressure distribution around the prementum of *Sympetrum*. Only pressure exceeding ambient pressure is displayed. It shows no high-pressure region at the opening which might complicate prey capturing. E) Vector field in the median plane showing flow speed distribution around the prementum. Inside the premental cavity, the flow speed is increasing and directed into the cavity, which correlates with regions of lower pressure. CFD Simulations based on kinematic data at timepoint 6 (0.44m/s flow velocity and α = 44°).

#### Experimental validation

We performed four additional high-speed recordings with particles (algae) in the fluid to visualize actual flow patterns (Fig. 10, supplementary video S9). The results are consistent with our simulation outputs. We see a high-pressure region below the prementum, which is indicated by outflow in the region, but no bow wave is forming in either species and particles are flowing past the palps over/into the prementum. Furthermore, algae are visibly retained by the setae (missing particles in figure 10) and some non-steady effects can be observed (supp. video S9). After the palps have closed, the PLM slows down, leading to a flow out of the premental cavity due to fluid inertia and a pronounced bow wave.

**Figure 10).**
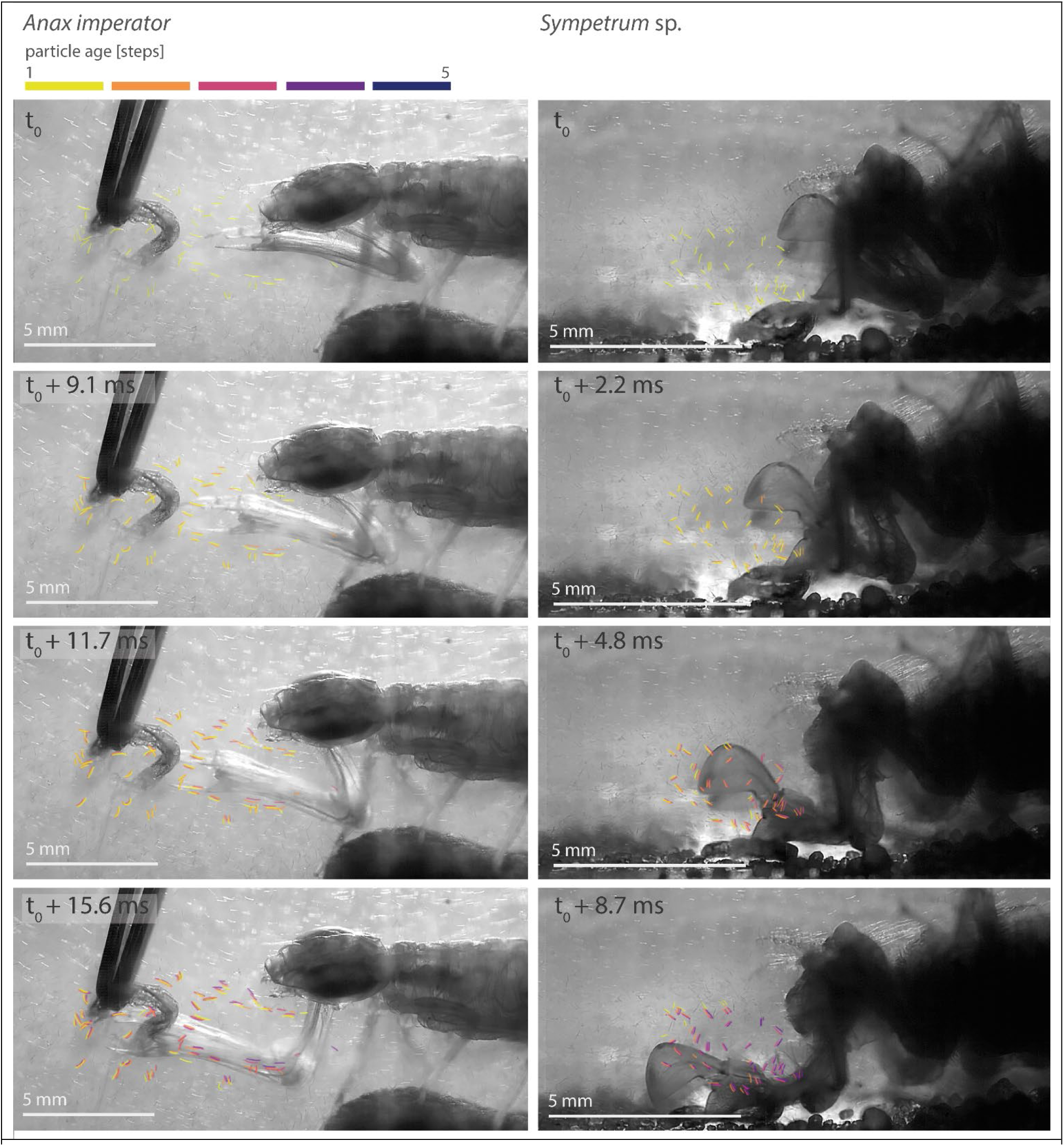
Motion of the fluid during a predatory strike. A) Still frames from HS-Video show characteristic points during a predatory strike of *Anax* (l) and *Sympetrum* (r). *Spirulina* algae were used as particles to observe the surrounding water. Characteristic particles are highlighted to visualize changes in particle position. Particle positions from previous images are shown in a different colour depending on their age, with the newest position in red. Both series show very little bow wave formation, as particles move barely compared to the motion of the PLM. Especially in *Anax*, we can also observe motion against the direction of the PRM above it (indicating a low-pressure zone), while particles below the PRM are pushed down. The corresponding videos can be found in supplementary material S9.

## Discussion

### Kinematics of the predatory strike

We derived translational and rotational velocity, in an attempt to approximate the flow speed at the tip of the prementum as closely as possible. Our results showed no significant difference between both species.

We assessed the angle of attack (*α*) of the prementum for the first time. *α* is a typical characteristic of airfoils and defined as the angle between the chord line of the airfoil, and the direction of flow (e.g. Gracey 1958). In a more general way, *α* can be understood as the orientation of a streamlined object towards the direction of flow. In both species, we see a decrease in median α as velocity increases. Yet we found a higher α in *Sympetrum*. The trend is visible over all timepoints (Fig. 2) and the difference is statistically significant at three timepoints. In general, a higher α leads to an increase in the area exposed to the flow, and drag (and lift) coefficients increase (Fig. 6) (Hoerner 1965; Gur et al. 2010).

### Characteristics of drag

The most relevant factors responsible for the amount of drag are the shape of an object, its size and orientation, as well as the velocity of the surrounding water. The drag/lift curves (Fig. 3) roughly reflect the velocity profiles (Fig. 2 B). Yet velocity changes alone do not completely explain the observed results (Fig. 3): for example, we find the maximum amount of drag and lift in *Sympetrum* at timepoint 5, whereas the maximum velocity occurs at timepoint 6. This can be explained by *α* changing over time. Changing the orientation changes flow patterns around the object (given that the object is not a sphere), which changes both the amount of drag, as well as the drag coefficient (Hoerner, 1965; Vogel, 1997). At timepoint 4, *α* is higher, leading to a higher amount of drag.

To assess the relationship between drag and *α*, we tested different values of *α* (0°-70°) at the same velocity (Fig. 6). The drag coefficients are in a range that would be expected for a flat, streamlined body (Hoerner 1965; Vogel 1997). In *Anax*, it increases proportionally with *α* as more and more surface area is exposed to the flow (Fig. 6). The prementum of *Sympetrum* shows a different pattern. The drag coefficient is minimal at an *α* of 30°. In general, the drag coefficient of the prementum of *Sympetrum* is less responsive to changes of *α* as in *Anax* (Fig. 6). This corresponds to the observation of a higher *α* during the predatory strike in *Sympetrum* (Fig. 2). We assume that these two different minima are due to the different shapes of the prementum. If we look at longitudinal sections of the prementum (Supplementary figure S4 A, B), the prementum of *Anax* is roughly represents a straight slender shape. Yet in *Sympetrum* the anterior part (ligula) is angled downwards. Hence it is more parallel to the flow at higher *α*. According to the simulation, we would expect an *α* of around 10° in *Anax*, when strike velocity is high, and around 30° in *Sympetrum*. However, our kinematic data (Fig. 2) shows that the measured *α* is higher in both species in vivo, hence more factors are to be considered.

Both prementa generate substantial amounts of dynamic lift. According to the principle of Bernoulli/Venturi, lift is the result of a pressure difference between the upper and lower side of an object (Hoerner, 1965; Vogel, 1997). This occurs in both prementa (Figs. 4, 7) and the amount of lift is dependent on *α* (Fig. 6). The prementum of *Sympetrum* requires a higher *α* to achieve the same lift coefficient as *Anax* but surpasses the lift coefficient of *Anax* at high *α*. For both species, lift generation is rather low for values of *α*, where the drag coefficient is minimal. Higher *α* values, observed during the strike, could thereby be explained by the necessity to generate lift, which is rather counterintuitive. One would expect the prementum to generate as little lift as possible. Either positive, as well as negative lift, would result in vertical motion of the prementum relative to the direction of travel, likely complicating prey targeting and also requiring additional energy. The amount of lift generated, however, is substantial and the calculated lift forces are almost as high as the drag force (Fig. 3). We hypothesise that this is to avoid the formation of a bow wave in front of the projecting prey capturing device, which may displace the prey item (Lauder and Prendergast 1992; Van Damme and Aerts 1997; Alfaro 2002). This behaviour would serve an analogous function to the compensatory suction feeding observed in vertebrates (Lauder and Prendergast 1992; Van Damme and Aerts 1997; Alfaro 2002; Holzman and Wainwright 2009), yet by deflecting the flow instead of expanding a cavity.

The findings are concurrent with the contours of pressure in figures 4 and 7. As the prey items, *Sympetrum* feeds on, tend to be rather small (especially compared to the usual prey of aeshnids) (Pritchard 1964), countering bow wave formation might be more important for capturing smaller prey, which would explain the more pronounced pressure difference between premental cavity and surrounding fluid.

The prementum of *Anax* is about three times longer than the prementum of *Sympetrum*. To compare these two shapes, we calculated the dimensionless drag coefficient (C_dw_) (formula 2, see Materials and Methods). C_dw_ only depends on the shape and orientation of the object (as long as the density of the medium does not change and the Reynolds numbers are comparable). The Reynolds number of both movements is within a similar order of magnitude (^~^3700 in *Anax* vs ^~^1800 in *Sympetrum*). These values are comparable to the Reynolds numbers occurring in the flight of larger insects (Vogel 1997; Goyens et al. 2015). We consider the flow sufficiently similar to make the previous comparisons. Yet both drag and lift coefficients are functions of the Reynolds number (Hoerner 1965; Vogel 1997). When changing the size of the prementum, the drag and lift coefficients increase (supplementary figure S7). The increase of the drag coefficient in *Anax* might be explained by the fact that the streamlining is less effective at the smaller scale because the amount of viscous drag is higher (Vogel 1997). Vice versa, the less streamlined prementum of *Sympetrum* has a higher impact at the larger scale, which causes the drag coefficient to increase as well. In conclusion, the PLM might be tuned towards its respective flow regime. Comparing the PLM from different larval instars might be an interesting approach to test these hypotheses in future research.

Finally, we can combine the information on setae distribution with the result of our CFD simulations (Fig. 9). Our SEM images, which are concordant with earlier studies on libellulids (Tillyard 1917; Olesen 1979; Gupta et al. 1992) show that setae are covering the top of the premental cavity. With open labial palps, not the entire area over the prementum is covered, which would be anticipated for an efficient capturing device. Considering the streamlines inside the premental cavity (Fig. 9), the low-pressure area at the prementum might also serve to deflect water with prey items deeper into the premental cavity, where they are trapped. Once the labial palps close, the premental cavity is completely covered by the seta arrays on the prementum and labial palps and prey cannot escape (Fig 9B).

### Limitations of the model & Validation

In this study, we used several simplifications/approximations which must be addressed. First of all, we used a quasi-dynamic approach, to replicate the flow patterns during the actual predatory strike with a series of stationary simulations. This is likely to underestimate the total amount of drag: due to the added mass of fluid that sticks to the prementum due to viscous forces (Van Wassenbergh et al. 2008). Additionally, we neglected the rotational component of the motion of the prementum but took an approach using the sum of translational and rotational velocity at the tip of the prementum. This is likely to overestimate the actual drag, as the base of the prementum is moving slower than the tip. Yet we were not interested in calculating the actual drag with high precision, but rather in estimating the impact of the PLM shape on prey capturing. Therefore, we were most interested in fluid motion at the tip of the prementum, the point of contact between the predator and prey, which is simulated with the highest accuracy and calculating *α* for each timeframe ensured that the direction of flow is correct. Finally, it is important to note that we did not consider the drag generated by the postmentum, to reduce simulation complexity, as it is also located downstream from the prementum and the prey item. These aspects have to be kept in mind when interpreting the results.

To verify the predicted flow patterns, we observed predatory strikes with algae as flow particles (Fig. 10, supplementary video S9). The results corroborate our simulations concerning bow wave prevention. Outflow indicates a high-pressure region below the prementum and we observed fluid passing over/into the prementum without obstruction. Furthermore, these findings suggest that inertial effects of the fluid play little role in the prey capture event. We only see the ejection of fluid from the cavity due to deceleration of the PRM after the labial palps have closed around the prey, indicating the deceleration effects play little role in prey capturing (supplementary video 9). However, to fully analyse the flow field around the PLM, more advanced and expensive methods, like PIV or MRI flow measurements, would be necessary for future projects.

## Conclusion

We showed that both prementum types (in *Anax* and *Sympetrum*) are well streamlined and generate a low-pressure zone, likely to mitigate the formation of a high-pressure area in front of the prehensile labial mask, a process analogous in function to compensatory suction feeding. We also verified this hypothesis experimentally, showing there is no visible bow wave formation. Furthermore, the prementum of *Sympetrum* is optimised towards higher angles of attack, possibly due to the angled ligula. Both prementa are tuned towards their specific Reynolds number, which raises the question: How does the shape of the labium and flow patterns change over different sizes like in larval development? Finally, we found that the prementum of *Sympetrum* deflects the water flow above the prementum downwards, likely to push prey items into the setae-armoured trap located on the prementum. Both prementa function in a similar way, despite their different shapes. The deep premental cavity is likely an innovation to trap prey, and possibly to further stabilise the low-pressure region above the prementum. Our study explores the fluid mechanics of prey capturing in odonate larvae for the first time and explains how fluid mechanics might facilitate prey capturing in these aquatic predators. These findings also show an example of convergent evolution, as this bow wave mitigation strategy represents a functional analogue to compensatory suction feeding known from vertebrates, like turtles, salamanders, and fish.

Furthermore, we show, how small scale technical underwater grasping devices for differently sized objects may function, which could be an interesting starting point for biomimetic applications.

## Abbreviations

α: angle of attack
ρ: density
μ: dynamic viscosity
C_dw_: drag coefficient
C_lw_: lift coefficient
F_d_: drag force
F_l_: lift force
PLM: prehensile labial mask
R: Reynolds number
v: Velocity

## Ethics

No permission of research ethics or animal ethics committees was necessary. All applicable regulations concerning the protection of free-living species were followed

## Data accessibility

Raw data can be obtained from the corresponding author upon request.

## Authors’ contributions

AK, MB, SNG and SB designed the project and developed the concept of the study. AK and SB performed the high-speed video recordings and analysed the footage. AK and MB designed the setup for the CFD simulation and evaluated the results. AK performed the CFD simulations and SEM observations. SB performed CT scanning of the specimens. AK wrote the manuscript. All authors read and edited the manuscript, agreed to the content therein and approved the final version of the manuscript.

## Acknowledgements

Finally, we would like to thank everyone, who supported us with this study: T. Büscher and H. Tramsen for support with statistics and data processing. B. Josten and H.-L. Troeger for support with the high-speed video experiments. Furthermore, we would like to thank S. Bodenstein, A. Georg, H. Jeske and K. Lehmann for providing dragonfly larvae for the study, O. Mudimo for providing *Spirulina*, and L. Hindenberg & H. Poggemann for proofreading. All members of the Functional Morphology and Biomechanics group at Zoological Institute of Kiel University are also greatly acknowledged for their continuous support.

## Supplementary Material

**Supplementary Figure S1).**
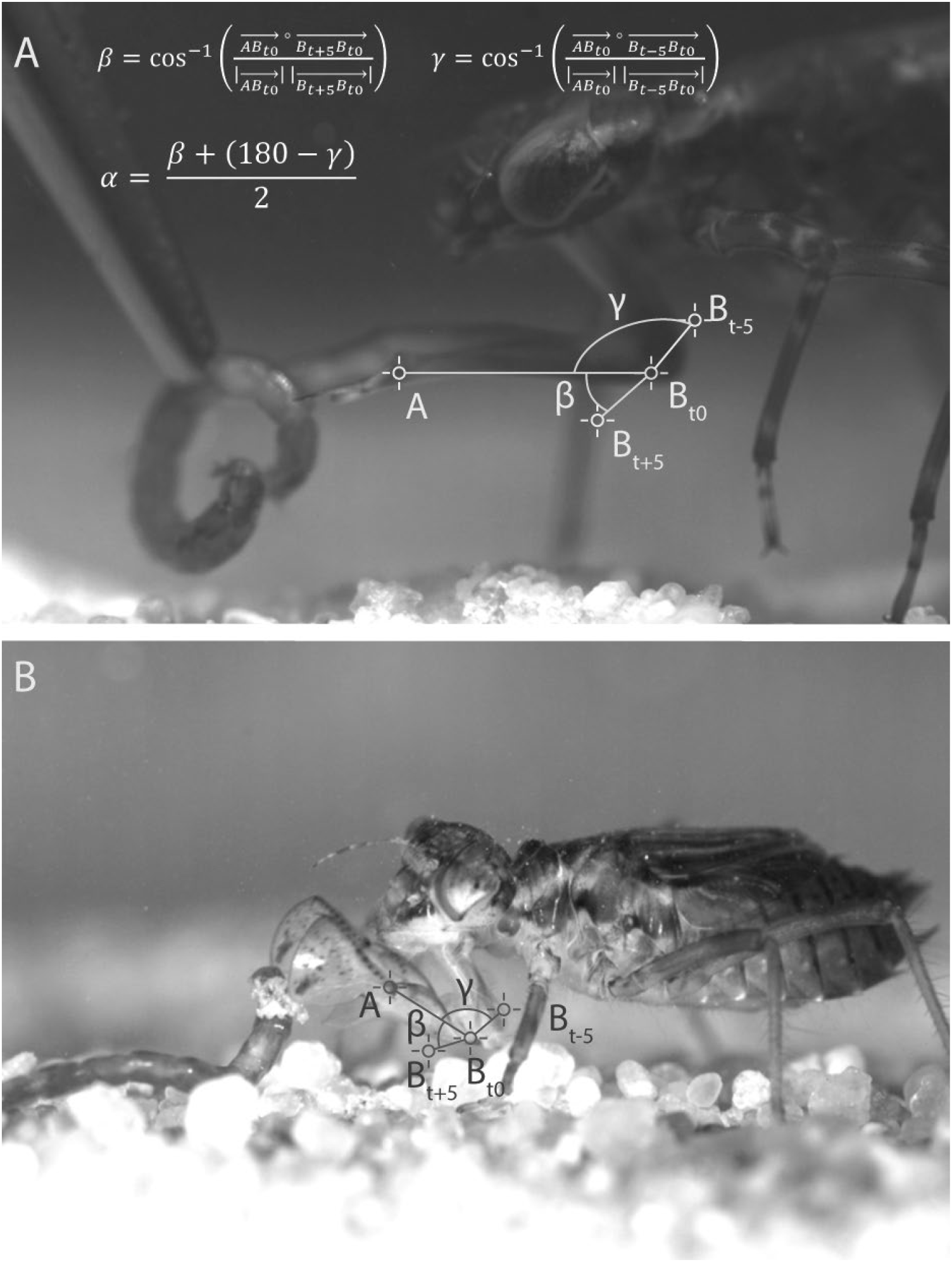
Exemplary still frames from two High-speed Videos showing the tracking points and calculated vectors. A) *Anax imperator* B) *Sympetrum* sp. Videos were recorded at 5400 fps. The formula for α calculation is shown in A), with B_t0_ being the location of the tracking point in the frame for which α is calculated, B_t+5_ being the location of the tracking point 5 frames later and B_t-5_ the location 5 frames earlier.

**Supplementary Table S2).** Calculation of Reynolds Number and C_dw_ and C_lw_ for one exemplary timepoint.

S2) Supplementary Table
| **R** | **$\rho$ [kg/m<sup>3</sup>]** | **l [mm]** | **v [ms<sup>-1</sup>]** | **$\mu$ [Pa\*s]** | **R** |
| --- | --- | --- | --- | --- | --- |
| *Anax* | 997.04 | 6.75 | 0.49 | 891 | 3701 |
| *Sympetrum* | 997.04 | 3.6 | 0.44 | 891 | 1773 |
| **C<sub>dw</sub> (timepoint 6)** | **$\rho$ [kg/m<sup>3</sup>]** | **A<sub>w</sub> [cm<sup>2</sup>]** | **v [ms<sup>-1</sup>]** | **F<sub>d</sub> [mN]** | **C<sub>dw</sub>** |
| *Anax* | 997.04 | 2.084 | 0.49 | 1.261 | 0.0051 |
| *Sympetrum* | 997.04 | 0.499 | 0.44 | 0.501 | 0.0106 |
| **C<sub>lw</sub> (timepoint 6)** | **$\rho$ [kg/m<sup>3</sup>]** | **A<sub>w</sub> [cm<sup>2</sup>]** | **v [ms<sup>-1</sup>]** | **F<sub>l</sub> [mN]** | **C<sub>lw</sub>** |
| *Anax* | 997.04 | 2.084 | 0.49 | 1.377 | 0.0552 |
| *Sympetrum* | 997.04 | 0.499 | 0.44 | 0.417 | 0.0865 |

**Supplementary Table S3).** Exemplary results of mesh analysis for *Sympetrum*.

S3) Supplementary Table
| Parameter | Final Resolution | 2x Resolution | Ratio |
| --- | --- | --- | --- |
| No. of elements | 4,681,683 | 18,230,392 | 25.68 |
| F<sub>drag</sub> [N] | -0.000510598 | -0.000514853 | 99.17 |
| F<sub>lift</sub> [N] | 0.0004257 | 0.000417959 | 98.18 |

**Supplementary Figure S4).**
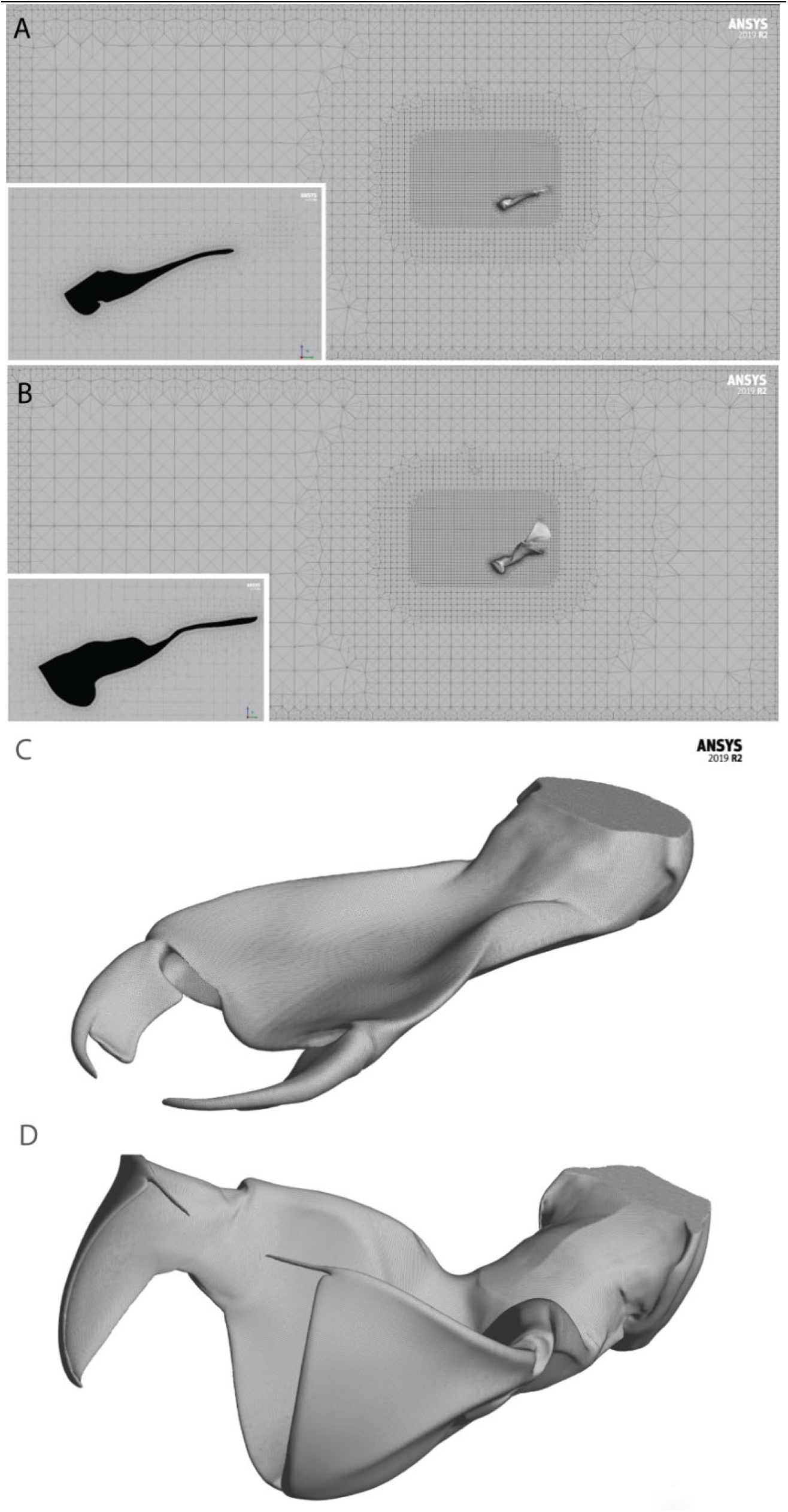
3D meshes used for CFD simulations. A, B) Exemplary meshes for CFD Simulation as generated by ANSYS ICEM. A) *Anax imperator*, B), *Sympetrum* sp. Large images show a longitudinal section through the entire domain, insets provide a close-up around the model to show prism layers. C.D) 3D models of the prementum based on CT scans of the head of a specimen of each *Anax imperator* C) and *Sympetrum* sp. D). The models were retopologised to ensure a clean topology without holes or intersections. The wireframe shows surface mesh resolution.

**Supplementary Figure S5).**
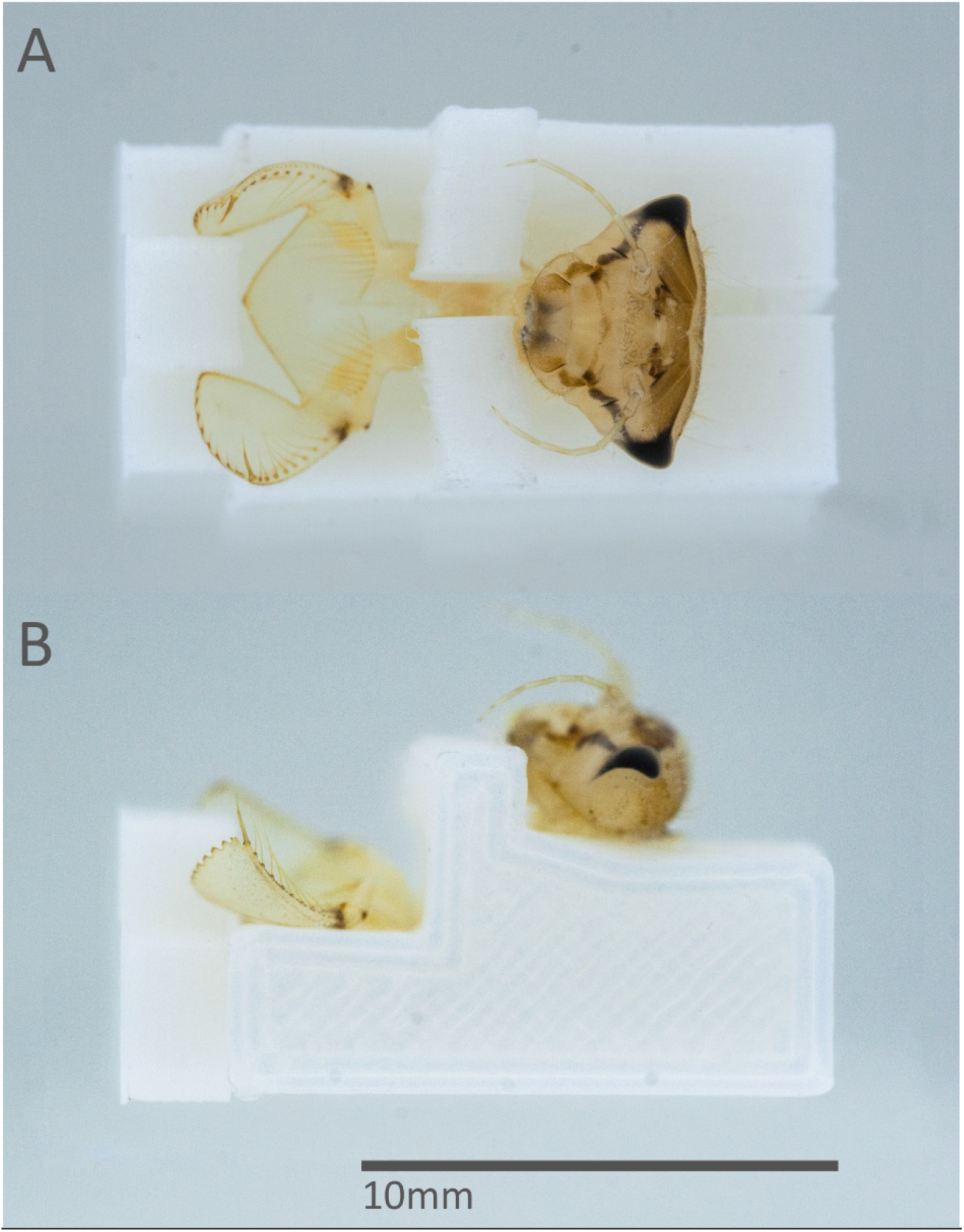
Expendable 3D printed brace for critical point drying of the prehensile labial mask in an opened state. The brace was designed in Blender and printed on a Prusa i3 Mk 3 FDM Printer using PLA Filament.

**Supplementary Figure S6).**
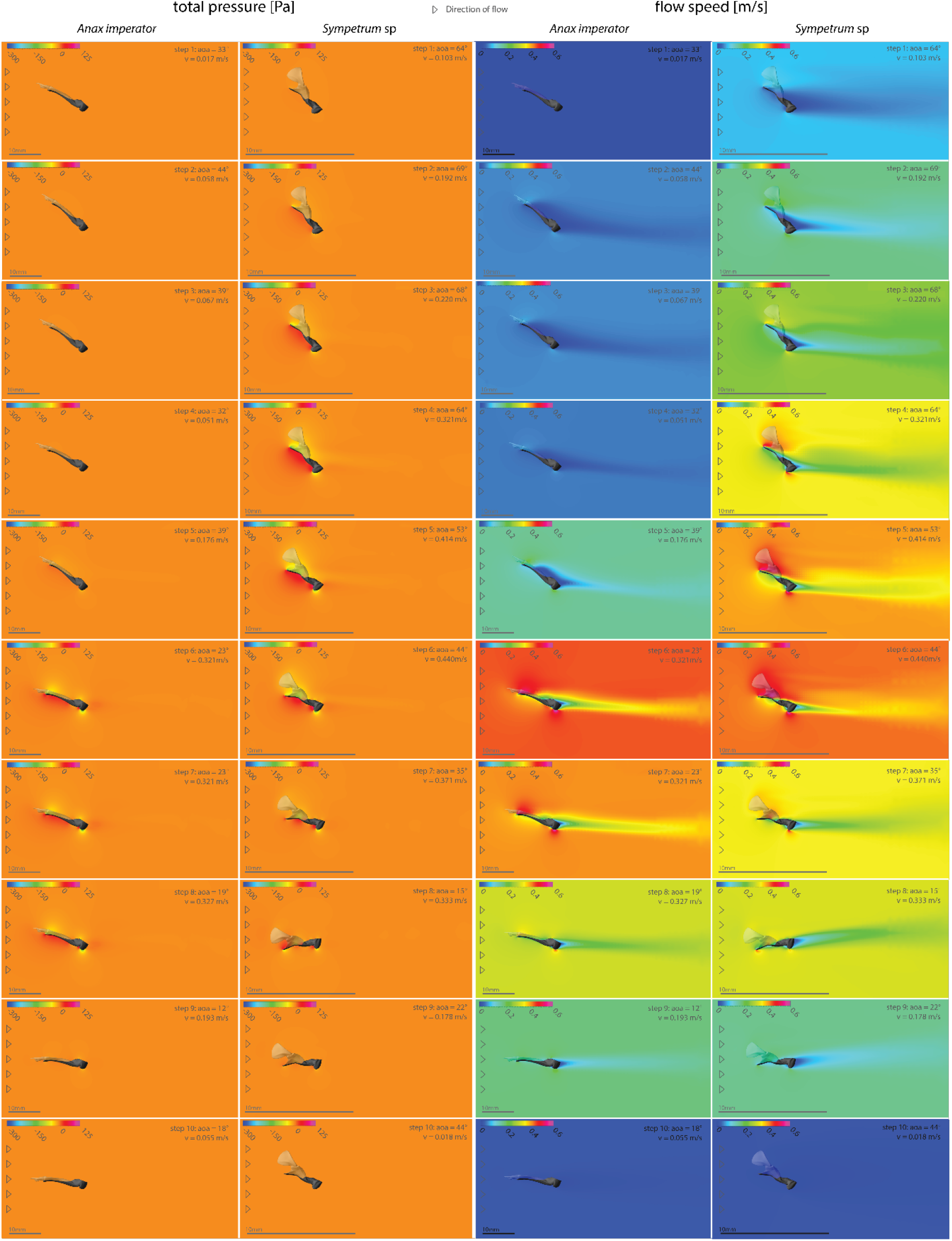
Flow characteristics change during the predatory strike. Contours of pressure (left) and contours of total flow speed (right) around the median line of the prementum of *Anax* l) and *Sympetrum* r) at all timepoints. l) Colour indicates fluid pressure around the prehensile labial mask. Note the formation of low-pressure regions on the dorsal side of both labial masks (yellow areas). r) Colour indicates total flow speed around the prehensile labial mask (the colour range is identical in all images). Note the flow separation at higher α in *Anax* (Step 3,5). This does not occur in *Sympetrum*. CFD Simulations were based on kinematic data from HS Videos (Figure 2).

**Supplementary Figure S7).**
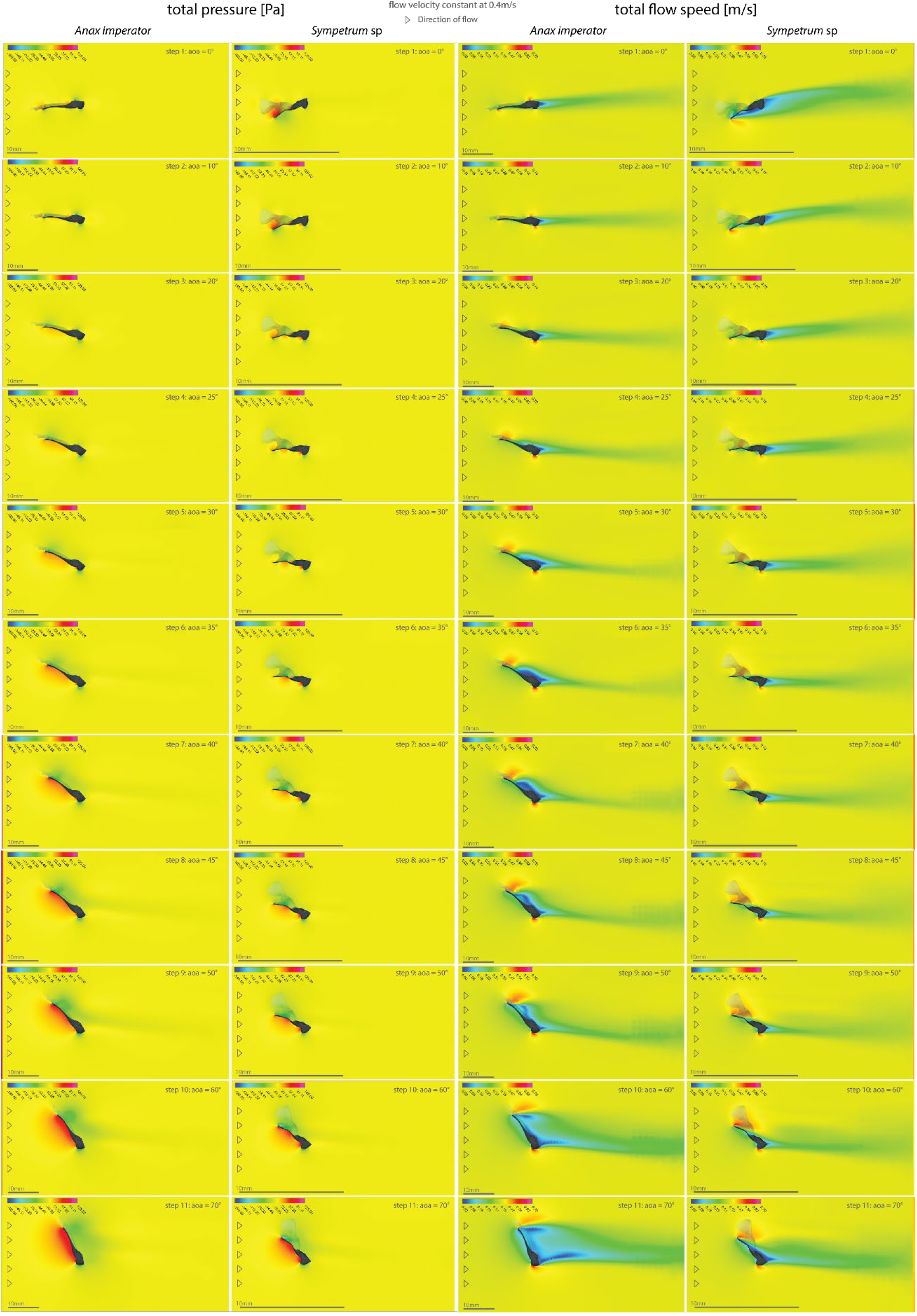
Flow characteristics change with the angle of attack. Left) Contours of pressure around the median line of the prementum of *Anax* l) and *Sympetrum* r). Colour indicates fluid pressure around the prehensile labial mask. Note the formation of low-pressure regions on the dorsal side of both labial masks at higher α. In *Sympetrum* a high-pressure area forms in the premental cavity at low α (e.g. 10°); right) Contours of total flow speed around the median line of the prementum of *Anax* l) and *Sympetrum* r). Colour indicates total flow speed around the prehensile labial mask (the colour range is set to a local range, to enhance pattern visibility). Note the separation of flow occurring in the prementum of *Anax* at higher α. This does not occur in *Sympetrum*. CFD Simulations were run at 0.4m/s fluid velocity and α was varied between 0° and 70° (Figure 6).

**Supplementary Figure S8).**
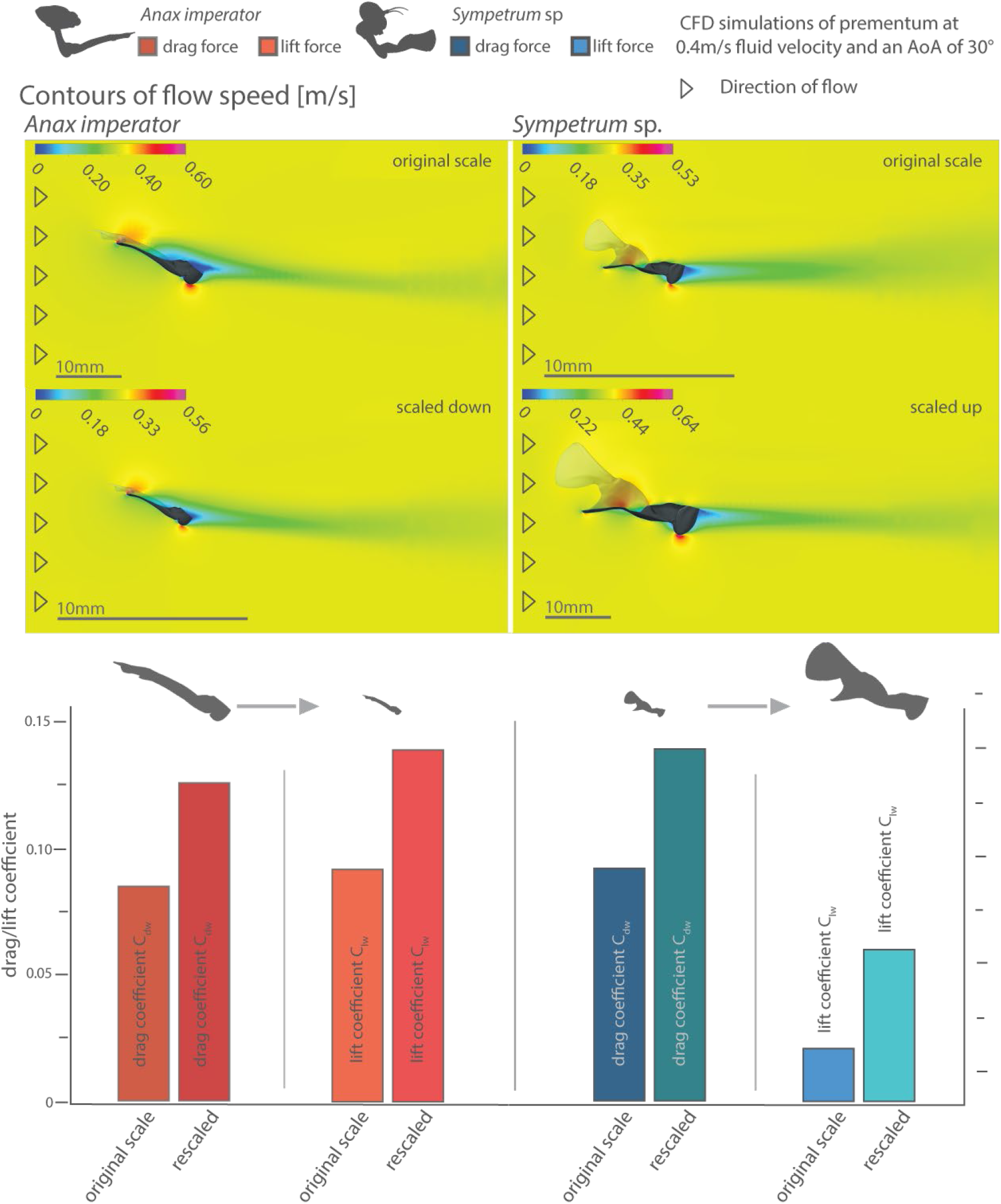
Changing the size increases drag and lift coefficient. Scaling the prementum of *Anax* down to the size of the prementum of *Sympetrum* increases its drag and lift coefficient. Scaling the prementum of *Sympetrum* up also increases its drag and lift coefficient. The overall flow patterns are similar. We only see a lower degree of flow separation in the downscaled *Anax* prementum due to the lower Reynolds number.

### S9) Supplementary Video

Video of predatory strikes with visualised flow.

